# Epigenome-wide change and variation in DNA methylation from birth to late adolescence

**DOI:** 10.1101/2020.06.09.142620

**Authors:** Rosa H. Mulder, Alexander Neumann, Charlotte A. M. Cecil, Esther Walton, Lotte C. Houtepen, Andrew J. Simpkin, Jolien Rijlaarsdam, Bastiaan T. Heijmans, Tom R. Gaunt, Janine F. Felix, Vincent W. V. Jaddoe, Marian J. Bakermans-Kranenburg, Henning Tiemeier, Caroline L. Relton, Marinus H. van IJzendoorn, Matthew Suderman

**Affiliations:** Department of Child and Adolescent Psychiatry/Psychology, Erasmus MC, University Medical Center Rotterdam, Rotterdam, the Netherlands; Generation R Study Group, Erasmus MC, University Medical Center Rotterdam, Rotterdam, the Netherlands; Institute of Education and Child Studies, Leiden University, Leiden, the Netherlands; Lady Davis Institute for Medical Research, Jewish General Hospital, Montreal, Qc, Canada; Department of Epidemiology, Erasmus MC, University Medical Center Rotterdam, Rotterdam, the Netherlands; Department of Psychology, Institute of Psychology, Psychiatry & Neuroscience, King’s College London, London, UK; MRC Integrative Epidemiology Unit, Population Health Sciences, Bristol Medical School, University of Bristol, Bristol, UK; Department of Psychology, University of Bath, Bath, UK; School of Mathematics, Statistics and Applied Mathematics, National University of Ireland, Galway, Ireland; Molecular Epidemiology, Department of Biomedical Data Sciences, Leiden University Medical Center, Leiden, The Netherlands; Department of Pediatrics, Erasmus MC, University Medical Center Rotterdam, Rotterdam, the Netherlands; Clinical Child & Family Studies, Vrije Universiteit Amsterdam, Amsterdam, the Netherlands; Department of Social and Behavioral Science, Harvard TH Chan School of Public Health, Boston USA; Department of Psychology, Education and Child Studies, Erasmus University Rotterdam, Rotterdam, the Netherlands; School of Clinical Medicine, University of Cambridge, Cambridge, UK

**Author notes:** both authors contributed equally to this manuscript (Alexander Neumann, Charlotte A. M. Cecil). Corresponding author: Matthew Suderman.

**Keywords:** EWAS, longitudinal, ageing, DNA methylation, change, childhood, trajectories, inter-individual variation, Generation R, ALSPAC

## Abstract

DNA methylation (DNAm) is known to play a pivotal role in childhood health and development, but a comprehensive characterization of genome-wide DNAm trajectories across this age period is currently lacking. We have therefore performed a series of epigenome-wide association studies in 5,019 blood samples collected at multiple time-points from birth to late adolescence from 2,348 participants of two large independent cohorts. DNAm profiles of autosomal CpG sites (CpGs) were generated using the Illumina Infinium HumanMethylation450 BeadChip. Change over time was widespread, observed at over one-half (53%) of CpGs. In most cases DNAm was decreasing (36% of CpGs). Inter-individual variation in linear trajectories was similarly widespread (27% of CpGs). Evidence for nonlinear change and inter-individual variation in nonlinear trajectories was somewhat less common (11% and 8% of CpGs, respectively). Very little inter-individual variation in change was explained by sex differences (0.4% of CpGs) even though sex-specific DNAm was observed at 5% of CpGs. DNAm trajectories were distributed non-randomly across the genome. For example, CpGs with decreasing DNAm were enriched in gene bodies and enhancers and were annotated to genes enriched in immune-developmental functions. By contrast, CpGs with increasing DNAm were enriched in promoter regions and annotated to genes enriched in neurodevelopmental functions. These findings depict a methylome undergoing widespread and often nonlinear change throughout childhood. They support a developmental role for DNA methylation that extends beyond birth into late adolescence and has implications for understanding life-long health and disease. DNAm trajectories can be visualized at http://epidelta.mrcieu.ac.uk.

## Introduction

DNA methylation (DNAm), an epigenetic process whereby DNA is modified by the addition of methyl groups, has gained increasing attention over the past few decades, due to its pivotal role in development. *In utero*, DNAm is involved in a range of essential processes including cell differentiation^1–3^, X-chromosome inactivation^4^ and fetal growth^5^. Its role extends well beyond birth, e.g. by maintaining cell type identity and genome stability^6–8^, responding to environmental exposures^9–11^, and its involvement in immune^12^ and neural development^13^. Since it is influenced by *both* genetic and environmental factors^14,15^, DNAm has also emerged as a key mechanism of interest for understanding the gene-environmental interplay in normal ageing and disease development.

Numerous studies have identified strong associations between DNAm and age. While most have relied on cross-sectional data^16–18^, but a few have utilized longitudinal measurements of DNAm within individuals^19–23^. Longitudinal measurements allow one to distinguish intra-individual change from inter-individual differences in change, thereby greatly improving the power to detect change over time and to identify differences between individuals^24^. Identifying and characterizing CpGs for which DNAm changes differently over time between individuals (i.e. inter-individual variation in change) is a necessary step in identifying genetic and environmental influences on the methylome as well as their potential impact on health outcomes^25^. Moreover, longitudinal designs facilitate the study of nonlinear trajectories^26,27^, which might help to identify sensitive periods for DNAm change in development. To date, the largest epigenome-wide longitudinal study on DNAm included 385 elderly individuals who were followed up to five times over a maximum period of 18 years, identifying DNAm change at 1,316 CpG (Cytosine-phosphate-Guanine) sites^19^ and inter-individual variation at change at 570 CpGs^20^. Yet, little is known about DNAm trajectories across early development, as existing studies in *childhood* DNAm typically have been limited by small sample sizes^21,23^, short time-periods^22,28^ or focused on specific CpGs in relation to maternal smoking^29^, birthweight^30^, or maternal BMI^31^.

In the current study, we aim to provide a benchmark of typical epigenome-wide age-related DNAm trajectories within individuals, spanning the first two decades of life. This study combines repeated measurements of DNAm at nearly half a million CpG sites across the genome from two large population-based cohorts, the Generation R Study and Avon Longitudinal Study of Parents and Children (ALSPAC), to form one integrated dataset with four time-points of measurement. In a series of three epigenome-wide mixed model analyses we study linear (Model 1), nonlinear (Model 2) and sex-related (Model 3) trajectories of change across development. Further, we aim to identify CpGs for which trajectories vary between individuals (Model 1 and 2). Results are interpreted in the context of CpG location and biological pathways. The key findings are discussed here, full results per CpG can be freely accessed and visualized at http://epidelta.mrcieu.ac.uk/ [*note to reviewers: this is a demo website for now*].

## Results

### Cross-cohort comparability

Sample characteristics of 1,399 Generation R participants (total DNAm samples=2,333) and of 949 ALSPAC participants (total DNAm samples=2,686; Figure 1) are provided in Supplementary Table 1. After the DNAm datasets of the two cohorts underwent joint functional normalization (see Supplementary Figure 1 for distributions of mean DNAm levels), within-cohort stability of DNAm at birth and 6 or 7 years (in Generation R and ALSPAC, respectively) was compared. Stability of DNAm at individual CpG sites (437,864 autosomal sites) was estimated in three ways: relative concordance using Spearman correlations between time points, absolute concordance using intraclass correlations between time points (children with data for both time points: *n* Generation R=476, *n* ALSPAC=826), and change over time using change estimates from a linear mixed model (Model 1, online Methods) applied within each cohort (children with data for at least one of the two time-points: *n* Generation R=1,394, *n* ALSPAC=944). Estimates of all stability measures for both cohorts are depicted in Figure 2. Next, agreement of these stability estimates between the two cohorts was estimated with the Spearman (ρ) or Pearson (*r*) correlation (depending on normality of the data) across all CpGs, between the datasets. The Spearman correlation of the relative concordance was ρ=0.62, the Pearson correlation of the absolute concordance was ρ=0.60, and the Pearson correlation of the change estimates was *r*=0.86, indicating strong agreement between datasets. Based on these results the two datasets were joined to form one set with four different time-points of DNAm (birth, age 6/7 years, 10 years, 17 years).

**Figure 1.**
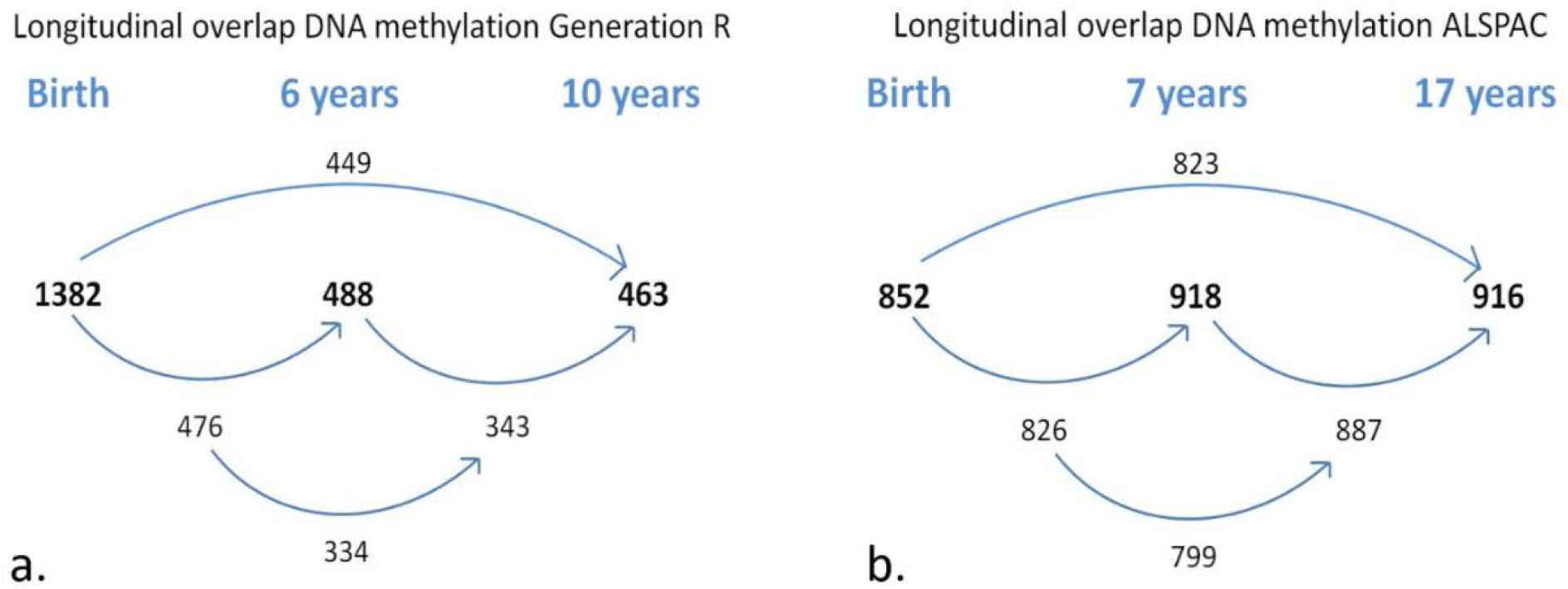
Longitudinal sample sizes. Sample sizes for (a) Generation R (N total children=1,399, N total DNAm samples=2,333); and (b) ALSPAC (N total children=949, N total DNAm samples=2,686). Bolded numbers represent total sample size at each time-point; non-bolded number refer to overlapping samples between time-points.

**Figure 2.**
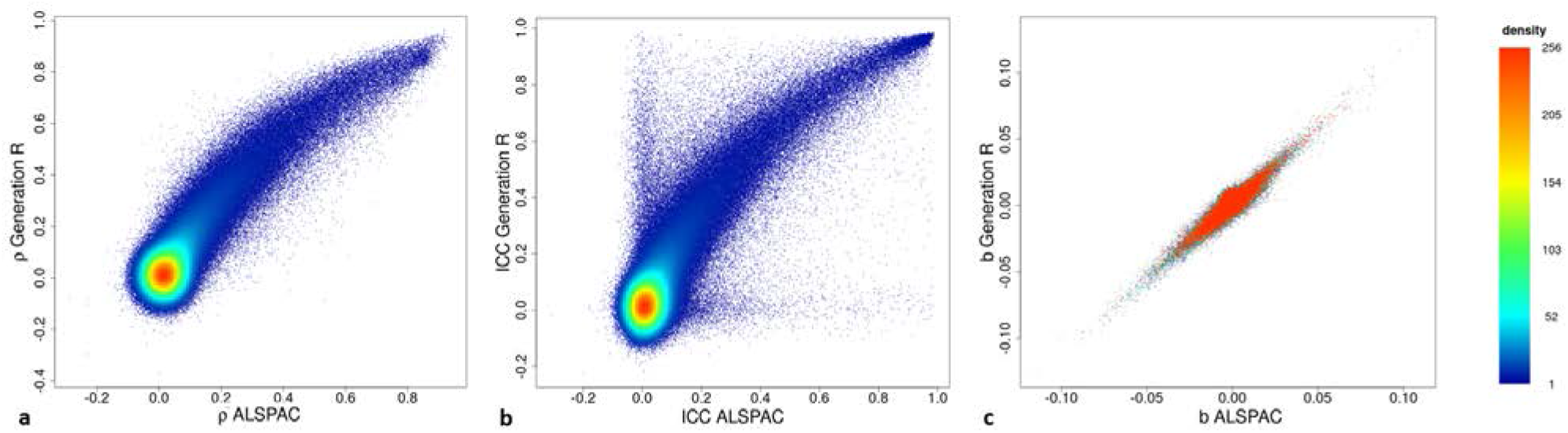
Scatterplots of within-cohort stability of DNA methylation. Showing (a) Spearman correlations, (b) intraclass correlation coefficients and (c) change estimates from birth to 6/7 years per CpG for Generation R and ALSPAC.

### Linear DNAm change from birth to early adulthood

Estimates of overall change in DNAm from birth to early adolescence (Model 1; see online Methods) indicated linear change at 51.6% of CpGs at a Bonferroni-corrected threshold (*P*<1×10^−07^) (Figure 3a and 3b). Specifically, DNAm decreased over time at 35.5% of all CpGs and increased at 16.0%. The mode intercept indicated that the decreasing CpGs were 88% methylated at birth (Figure 4). DNAm levels for increasing CpGs typically started at 5%.

**Figure 3.**
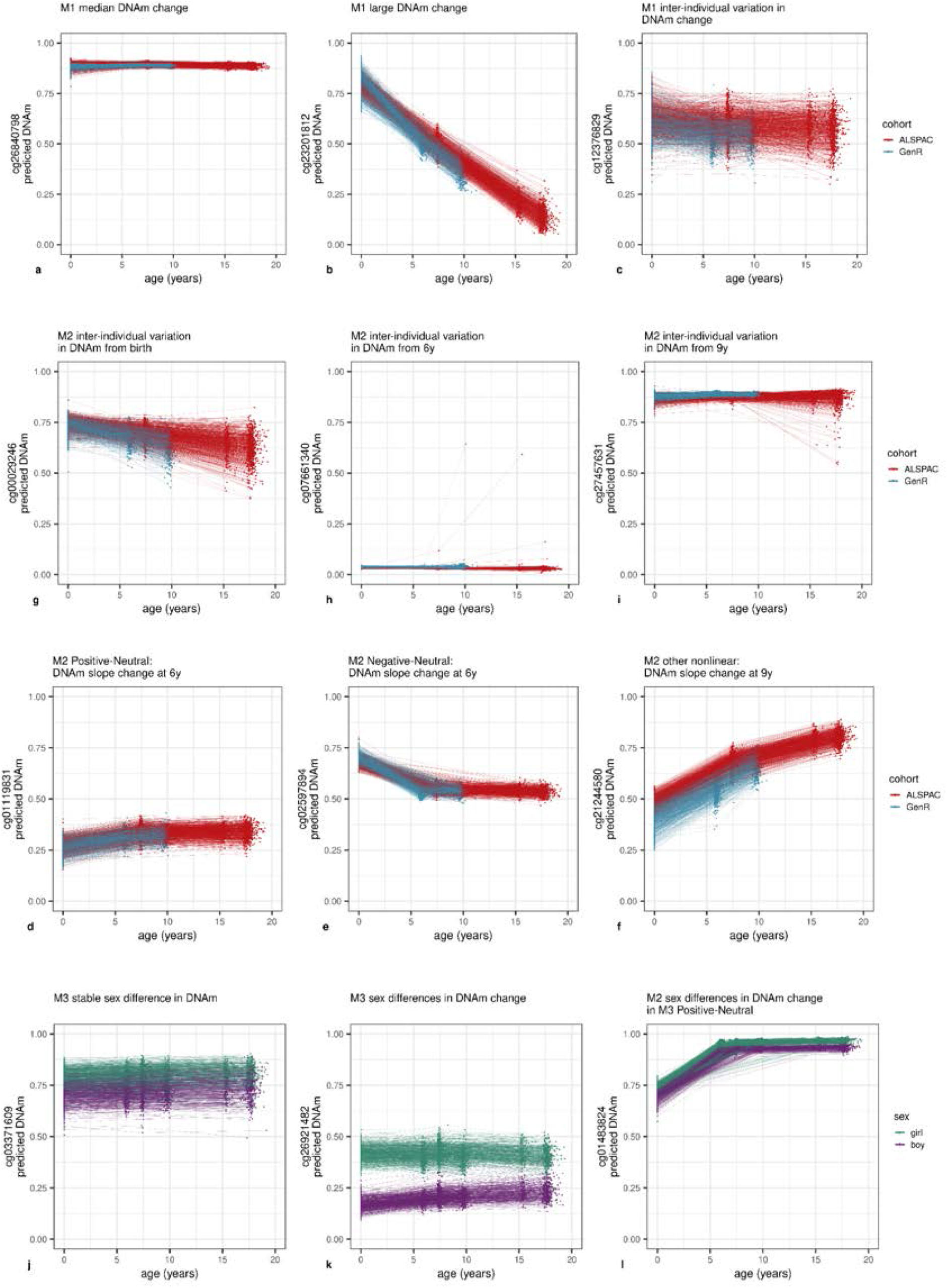
DNAm levels of selected CpG sites across childhood. Parts (a-c) show CpG sites with linear change over time (Model 1). A typical site is shown in (a), the site with the largest observed change in (b) and with inter-individual variation in DNAm change (c). Parts (d-f) show CpG sites with non-linear change (Model 2). A Positive-Neutral trajectory is shown in (d), a Negative-Neutral trajectory in (e) and a Positive-More Positive-Less Positive in (f). Parts (g-i) show CpG sites with inter-individual variation in change (Model 2). A site with slope variation from birth is shown in (g), slope change variation at 6 in (h) and slope change variation at 9 in (i). Parts (j-l) show CpG sites with sex-specific DNAm. A site with stable sex differences is shown in (Model 3) (j), sex-specific slope in (Model 3) (k) and sex-specific slope change at 6 in (Model 2) (l).

**Figure 4.**
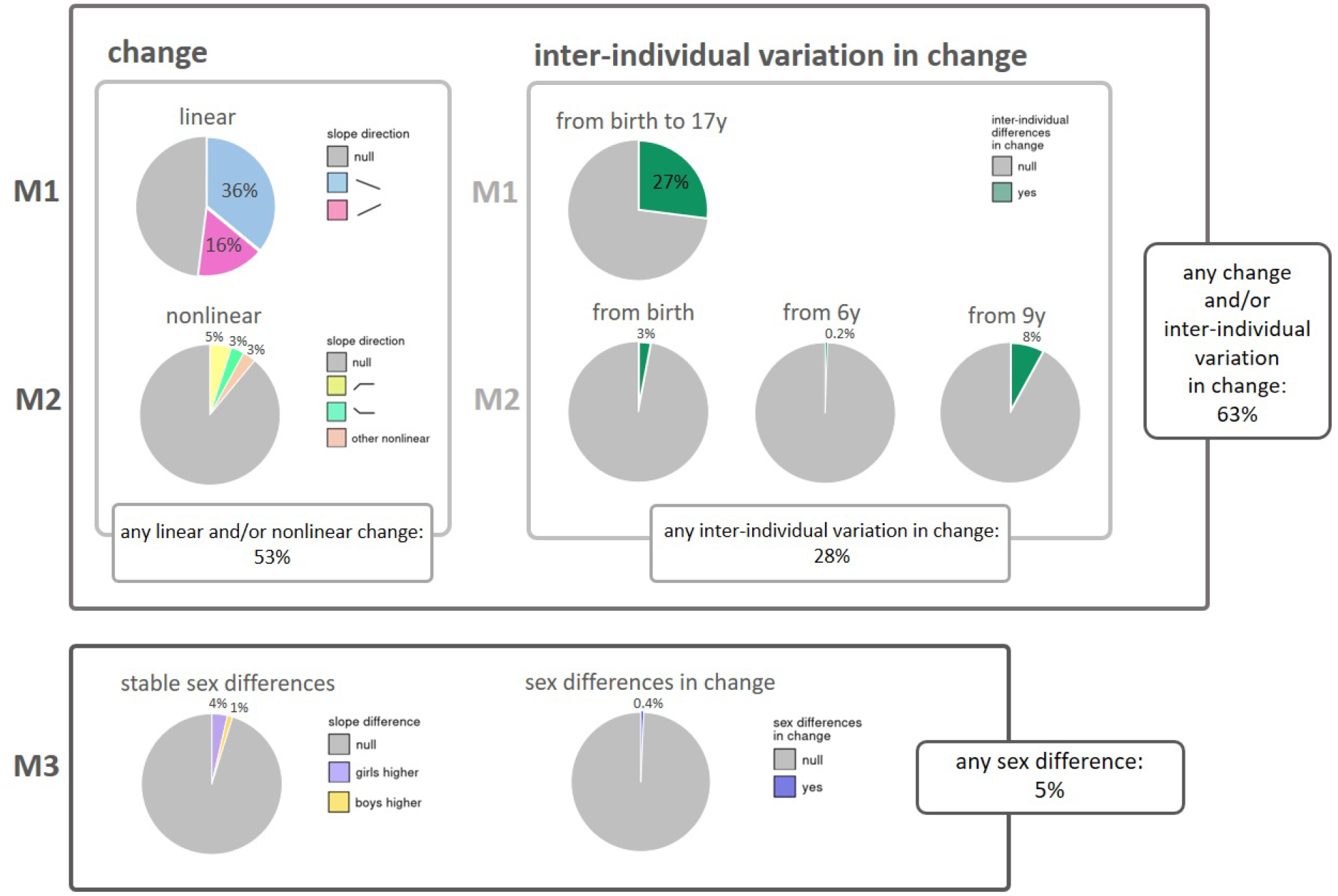
Overview of results from the three models. Model 1 (M1) was applied for overall change in DNA methylation and inter-individual variation in linear change; Model 2 (M2) for nonlinear change in DNA methylation and inter-individual variation in nonlinear change; and Model 3 (M3) for stable sex differences in DNA methylation and sex differences in change of DNA methylation (Sex by Time interaction). Percentages represent percentage of autosomal CpGs below Bonferroni-corrected threshold (P<1×10^−07^).

**Figure 5.**
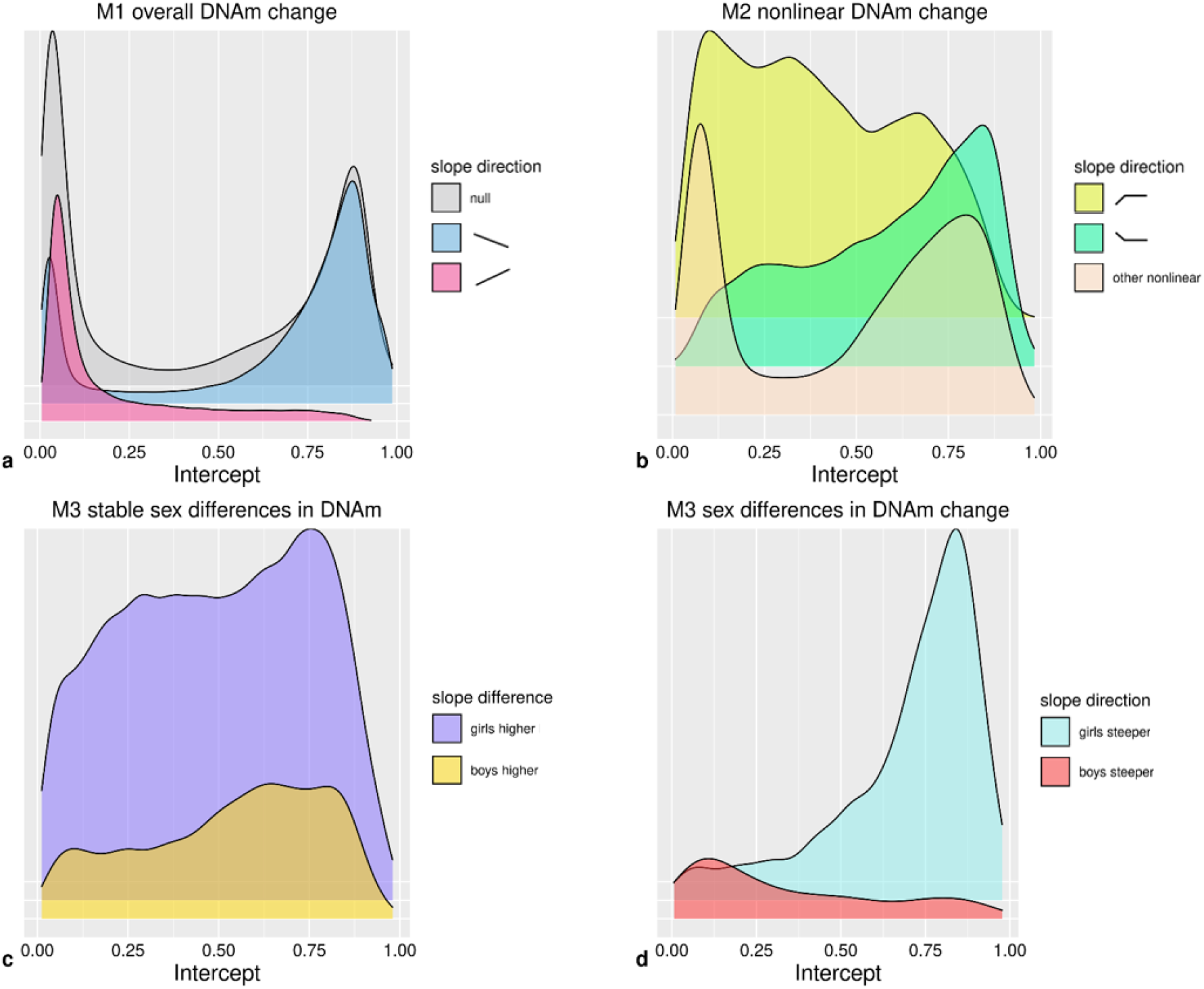
Density plots of intercepts of CpGs. Intercepts for CpGs with (a) directions of change in Model 1 (n=473,864); (b) nonlinear trajectories in Model 2 (n=52,043); (c) stable sex differences in Model 3 (n=22,821); (d) sex differences in DNAm change in Model 3 (n=1,768).

The mode estimate DNAm change was *b*=−9.24×10^−04^ (with corresponding mode *SE*=6.85×10^−05^), indicating an overall 0.09% DNAm decrease per year at a typical CpG site. This translates into a 1.66% decrease in DNAm over the course of 18 years. An example of a CpG site with a typical change in DNAm is depicted in Figure 3a. The largest observed absolute change in DNAm was *b*=−3.47×10^−02^ (*SE*=3.65×10^−04^, *P*<9.88×10^−324^), indicating an overall DNAm decrease of 62.5% over 18 years (Figure 3b). Only twenty-two CpGs showed an absolute change >50% over the course of 18 years (Supplementary Table 2). From this it follows that typically in (cord-/peripheral) blood tissue, DNAm levels for CpGs do not change from a fully unmethylated to fully methylated state, or vice versa, over the course of 18 years.

Further, we observed substantial inter-individual variation in linear DNAm changes over time at 27.4% of all CpGs (i.e. random slope variance was greater than zero at Bonferroni-corrected threshold *P*<1×10^−07^; Figure 3c). On average, this variation accounted for 2.7% (*SD*=1.5%) of all estimated inter-individual variation (for intercept, age, batch, and residual) at these CpGs. At 17.3% of all CpGs, we observed both change and inter-individual variation in change.

### Nonlinear DNAm change

Model 2 (see online Methods) was identical to Model 1 but permitted slope changes at ages 6 and 9 years to test for nonlinear DNAm trajectories. At 11.0% of CpGs a nonlinear trajectory was detected. Specifically, at 4.8% of all CpGs, DNAm increased from birth and remained stable from 6 onward (Positive-Neutral; Figure 3d). Second, at 3.1% of all CpGs, DNAm decreased from birth and then remained stable at 6 years (Negative-Neutral; Figure 3e). The remaining 3.0% of all CpGs followed other nonlinear trajectories (e.g. Figure 3f), with each trajectory observed in <1.0% of all CpGs. Overall, linear and/or nonlinear changes in Model 1 or 2 were observed in 52.6% of CpGs (Figure 3), indicating that most nonlinear patterns were also detected as linear patterns in Model 1.

Inter-individual differences in change (i.e. random variance in slopes) from birth onward was detected at 3.4% of all sites (Figure 3g), inter-individual differences in slope change at 6 years in 0.2% (Figure 3h), and inter-individual differences in slope change at 9 years at 8.2% of CpGs (Figure 3i). Inter-individual differences in slope (change) at each time-point were detected more often at CpGs with an increasing rather than decreasing overall DNAm change in Model 1 (*P*=2.37×10^−144^). Last, both Positive-Neutral and Negative-Neutral changes coincided more often with inter-individual variation from birth (*P*<9.88×10^−324^). Any inter-individual differences in change, detected by Model 1 or 2, was observed at 27.9% of CpGs. In total, Models 1 and 2 detected age-related change whether linear, non-linear or inter-individual differences in change at 62.8% of all CpG sites (Figure 3).

### Sex differences in longitudinal DNAm and DNAm change

According to Model 3 (online Methods), sex differences in DNAm were present at 4.9% of (autosomal) CpGs (Figure 3). Specifically, stable longitudinal sex differences (main sex effects) were observed at 4.8% of all (autosomal) CpGs (Figure 3j), and sex differences in DNAm change (sex by age interaction effects) were found at 0.4% of all (autosomal) CpGs (Figure 3k). At sites with stable sex differences, DNAm levels were higher in girls at 3.6% (Figure 3j) and lower at 1.2% of CpG sites. DNAm at sites with higher DNAm in girls tended to increase over time, whereas DNAm at sites with higher DNAm in boys tended to decrease (*P*=4.20×10^−205^). Most commonly (at 0.2% of all CpGs), DNAm was higher in girls at birth but DNAm in boys increased at a higher rate.

Both CpGs with stable sex differences and those with sex differences in DNAm change were less likely to show inter-individual variation than other sites (20.8% versus 27.5% and 18.1% versus 27.3%; *P*=5.36×10^−111^ and *P*=7.57×10^−18^). Finally, CpGs with stable sex differences or sex differences in DNAm change detected in Model 3 were much more likely to follow an overall Positive-Neutral trajectory of DNAm change detected in Model 2 than other CpG sites were (24.2% of CpGs with stable sex differences followed a Positive-Neutral trajectory versus 3.8% of other CpGs and 53.9% of CpGs with sex differences in DNAm change followed a Positive-Neutral trajectory versus 4.6% of other CpGs; *P*<9.88×10^−324^, *P*<9.88×10^−324^; Figure 3l). Albeit less prominently so, CpGs with stable sex differences or sex differences in DNAm change also more often followed a Negative-Neutral trajectory than other CpGs did (stable sex differences: 5.0% versus 3.0%, *P*=5.43×10^−62^; sex differences in DNAm change: 7.7% versus 3.1%, *P*<7.11×10^−28^).

### Follow-up analyses

Follow-up analyses were performed to understand how different types of age-related DNAm trajectories are distributed across the genome (Supplementary Tables 3-5). All reported enrichments have significance below a Bonferroni-corrected threshold of *P*<4.46×10^−04^, corrected for the number of chi-square tests (*n*=112). We further report enrichment of Gene Ontology (GO) pathways (nominal *P*<0.05) for genes annotated to CpG sites in each trajectory (Supplementary Tables 5-7). Last, we study enrichment of age-related DNAm trajectories in reported hits of different EWASs (Figure 6). All reported EWAS enrichments are below a Bonferroni-corrected threshold of *P*<2.16×10^−04^, corrected for the number of Fishers’ exact tests (*n*=231; Supplementary Table 8).

**Figure 6.**
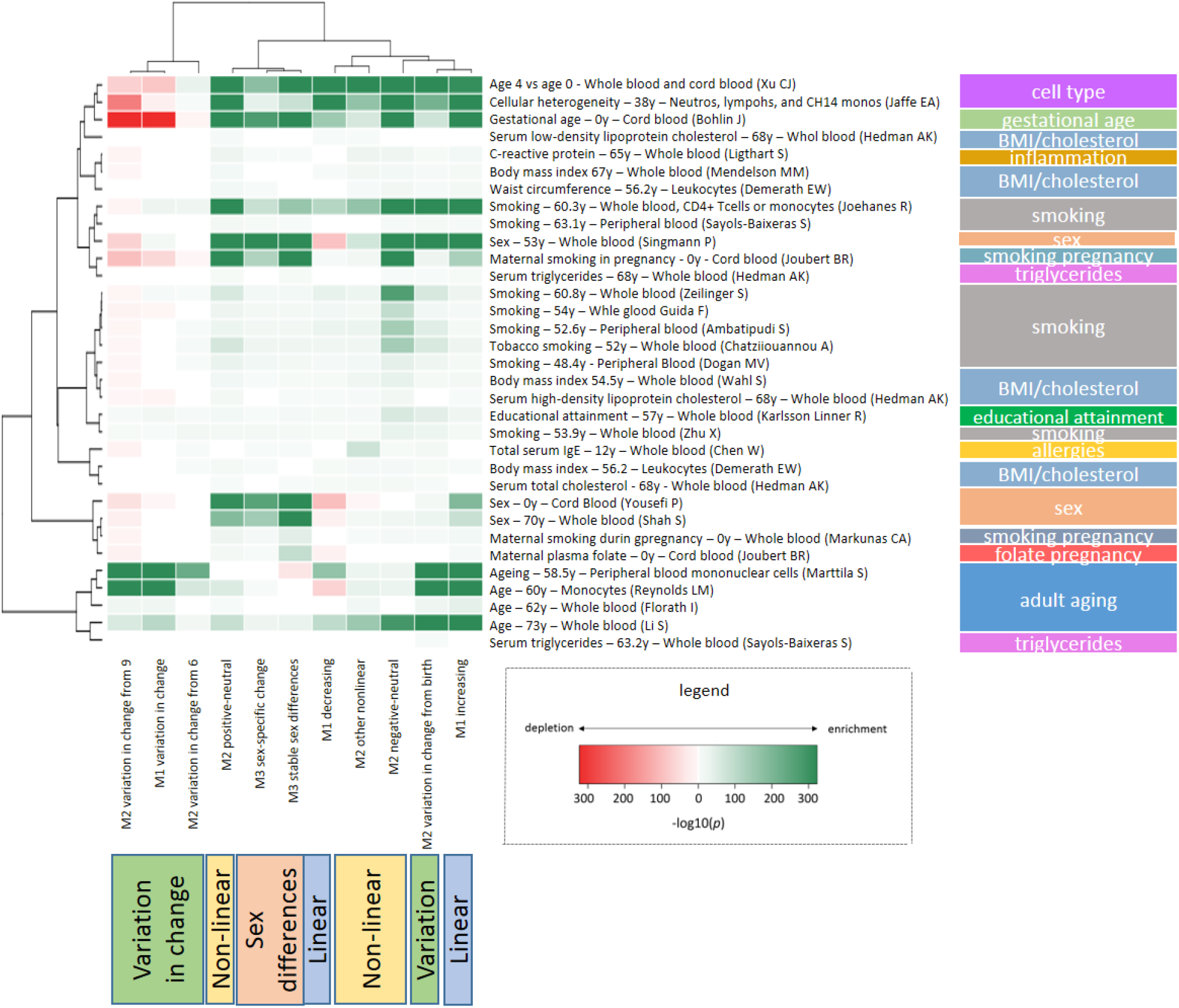
Enrichment of age-related trajectories in EWASs

#### Patterns of DNAm change and CpG location

CpG sites with DNAm change associated patterns were labeled by gene associated regions, CpG island associated regions, as well as enhancer elements. Although many exceptions exist, low levels of DNAm in the promoter area but high levels of DNAm in the gene body are generally associated with increased gene transcription^32,33^. CpGs annotated to TSS200 regions more often showed an overall DNAm increase (Model 1) than other CpGs (19.0% versus 15.6%), whereas CpGs annotated to the gene body more often showed an overall DNAm decrease than other sites (38.8% versus 33.7%). TSS200 CpGs showed less inter-individual variation in overall DNAm change than other sites (22.2% versus 28.1%), whereas gene body CpGs showed somewhat more inter-individual variation in overall DNAm change than other sites (28.9% versus 26.5%).

Promoter areas often coincide with CpG islands^34^. Here, 63.3% of TSS200 CpGs were also annotated to CpG islands. As in TSS200 areas, CpGs annotated to CpG islands had lower DNAm levels (mode M1 intercept 2.4% (*SD*=30.2%)), and more often showed an overall DNAm increase than other sites (25.2% versus 12.0%). DNAm sex differences were especially present in the shores of CpG islands compared to all other island associated regions (stable sex differences: 7.5% versus 4.0%, sex differences in DNAm change: 0.6% versus 0.3%).

Enhancers act on promoters to regulate gene transcription^35^. CpGs annotated to enhancer elements (2.0% of CpGs) tended to have low DNAm levels (mode M1 intercept 5.07%; *SD*=31.4%) and then increased with age more than other CpGs (23.9% versus 15.9%). Inter-individual variation in change from birth was more common at enhancer sites than at other sites (5.6% versus 3.3%).

#### Functional associations

Enrichment of Gene Ontology categories was tested for genes linked to CpGs with different DNAm trajectories. In short, genes annotated to CpGs with overall decreasing DNAm levels were enriched in immune-developmental functions, whereas those annotated to CpGs with increasing levels were enriched in neurodevelopmental functions. This pattern seemed even more pronounced at genes annotated to nonlinear Negative-Neutral and Positive-Neutral CpGs, with the former more often associated to immune-development and the latter to neurodevelopment. Genes linked to CpGs with stable sex differences and sex differences in DNAm change were enriched in pathways associated with sexual development, such as genital development, as well as pathways associated with neurodevelopment. Genes linked to CpGs with sex difference in DNAm change were also enriched in functions related to tooth and hair development.

#### Enrichment in EWASs

We further investigated functional relevance of CpG sites with age-related DNAm trajectories by testing enrichment with published EWAS associations (Figure 6)^28,36–61^. Unsupervised clustering of the enrichments shows that CpG sites with inter-individual variation in change over time have distinct enrichments and cluster differently from those with age-associated change that is consistent among individuals. The CpG sites of each age-associated DNAm trajectory were enriched with published age associations in adulthood. Multiple smoking EWAS clustered together with enrichment patterns exhibiting strongest enrichments among CpG sites with Negative-Neutral trajectories and mostly weak enrichments among CpG sites with inter-individual variation in change. Further, despite adjusting for cell count heterogeneity in our models, we observed enrichments of CpG sites that differ by white blood cell type among sites following nearly all age-associated trajectories. Finally, we observed enrichments of CpG sites associated with gestational age and prenatal smoking with sex-specific DNAm.

## Discussion

In this study we described changes in DNAm levels through the first two decades of human life. We examined DNAm levels per CpG by their linear association with age, their nonlinear trajectories and inter-individual variation in change, as well as sex differences and CpG characteristics. We found that about half of sites change: consistent linear and/or nonlinear DNAm change was found at 53% of sites. We further found that over a quarter of sites, 28%, were characterized by substantial inter-individual differences in the direction of this change. DNAm sex differences were present, but not abundant: 5% of autosomal sites displayed different DNAm levels or differences in change over time for girls and boys.

Specifically, we determined that DNAm at 52% of the measured methylome have some form of linear change from birth to late adolescence, with DNAm decreasing at 36% and increasing at 16% of CpGs. CpGs with decreasing DNAm tended to have high levels of DNAm and were more often located in gene bodies. CpGs with increasing levels of DNAm tended to have low levels of DNAm and were more likely to be located in promoter regions and at enhancers. The predominance of decreasing CpGs is in agreement with literature on epigenome-wide DNAm and age in cross-sectional research on children and adults^18,62^, as well as with longitudinal research in adults^19^.

Nonlinear DNAm trajectories were detected at 11% of CpGs, mostly involving changes in DNAm from birth to age 6 years, after which DNAm was more stable. We note that this could be due to cord blood being used to generate DNAm profiles at birth, whereas peripheral blood was used at later ages. A previous study^23^ including eight children showed that the cord blood DNAm profile at birth clustered separately from later peripheral profiles, after which DNAm changed gradually from 1, to 2.5, to 5 years. Such differences between DNAm in cord and peripheral blood might be due to uncaptured differences in white blood cell composition, as well as to different gene-regulatory functioning in the intra-uterine versus extra-uterine environment. On the other hand, Gene Ontology analyses showed that functional associations for positive and negative *linear* DNAm patterns, which are unlikely to be affected by tissue type, were consistent with functional associations for nonlinear positive and negative patterns, respectively (e.g. positive and negative up to 6 years, and then no change up to 18 years).

Specifically, sites with decreasing levels of DNAm, both with or without slope changes around the age of 6 years, were functionally enriched for immune-developmental pathways, and sites with increasing levels of DNAm, both with or without slope changes, were enriched for neurodevelopmental pathways. Since these observations were based on blood DNAm, it remains to be studied what roles genes linked to neurodevelopmental pathways play in in blood, or to what extent DNAm trajectories in blood mirror those in neural tissue.

Inter-individual differences in linear DNAm trajectories were found at 27% of CpGs, indicating change at different rates or directions for different individuals. Such sites tended to have overall increasing rather than decreasing levels of DNAm from birth to 18 years. This observation is consistent with the only other large study to examine inter-individual differences in DNAm change^20^, although we note that this study included elderly subjects and detected a smaller number of relevant CpGs. We are the first to investigate inter-individual differences in nonlinear DNAm trajectories. These were most often found in the slope change at 9 years (8% of CpG sites), indicating that most inter-individual differences in DNAm emerge after the first decade of life. More research is needed to understand if the direction of change in this period is determined by stimuli during that period, or rather by preceding, perhaps cumulative, exposures. However, it is clear that, given the high proportion of CpG sites with inter-individual variation in DNAm change over time that we have observed, it is important to restrict the range of ages of children included a single EWAS. Specific limits should be discussed given the rapidly growing number of studies generating DNAm profiles across childhood^63^.

Stable sex differences were found at 5% of autosomal CpGs, and sex differences in DNAm *change* were found at 0.4% of all CpGs. In general, if there were stable sex differences, girls had higher levels of DNAm (4% of all CpGs), in case of sex differences in DNAm change, boys had an accelerated upward change (0.2% of all CpGs). The direction of stable sex differences detected are congruent with a cross-sectional study on newborns, in which girls had higher DNAm levels than boys for the large majority of the 3031 significant autosomal CpGs^54^. Sex-discordant associations with age seemed to be more prevalent from birth to age 6 years than afterwards, suggesting that any phenotypic sex differences associated to DNAm would be established in early childhood. Their enrichment in the shores of CpG islands, areas at which DNAm has been associated with tissue differentiation and tissue-specific gene expression^64^, is consistent with the critical role that these processes play in sexual differentiation. Studies into sex differences in epigenetic regulation might want to focus on these locations.

We also found the other DNAm trajectories to be arranged throughout the genome in a nonrandom fashion. Earlier studies^32,65^ have shown that, for active genes, lower DNAm towards the promoter area (TSS200) and higher DNAm in the gene body relate to increased gene transcription. Here we add the observation that promoter DNAm tends to increase and gene body DNAm tends to decrease with age. From this finding, one might infer that a downregulation of gene expression takes place from birth to late adolescence. Enrichment analyses of published EWAS associations further showed that different traits and exposures exhibited distinct enrichment patterns among DNAm trajectories. For example, there were clear differences between smoking and BMI-related traits. Enrichment of sites with DNAm sex differences in EWASs on prenatal maternal smoking is consistent with studies finding that prenatal smoking affects traits such as birth weight^66^, brain development^67,68^, and attention^69^ differently in boys and girls. Clustering for prenatal maternal smoking EWASs also showed enrichment for CpGs with consistent change among individuals, not for CpGs with inter-individual variation in change. This may suggest a link with the well-known effects of prenatal smoking on childhood development since consistent DNAm change is more likely related to development or aging programming than inter-individual variation. This may explain why changes associated to prenatal smoking persist throughout life^70^. Notably, this pattern of change without inter-individual variation is visible in cg05575921, the *AHRR* CpG site strongly and persistently associated with prenatal smoking^71,72^ (Supplemental Figure 2; http://epidelta.mrcieu.ac.uk/).

‘Epigenetic age acceleration’ is a term coined to indicate the deviation of chronological age from age as estimated by an ‘epigenetic clock’ and is associated with disease risk and mortality^73^. Existing clocks are all linear models based on DNAm. Consequently, one might expect that all CpGs included in the clock model change linearly with age. Furthermore, to detect age acceleration, one would expect that these CpG sites would also vary between individuals. Surprisingly, many CpG sites included in the most popular clocks do not match these expectations^74,75^ (Supplemental Tables 9, 10). For example, we observe that over one-quarter and nearly one-half of the CpG sites included in the Horvath and Hannum clocks, respectively, follow non-linear DNAm trajectories in childhood. Given the widespread use of clocks to investigate biological aging, further investigation is warranted to better understand how, and perhaps if, associations using these clocks should be interpreted in child DNAm profiles.

We note three main limitations of our findings. First, the use of different tissue types (cord blood and peripheral blood) could account for some of the differences between birth and later time points, e.g. sites that increased or decreased between birth and 6, but did not show change after that. Generation of DNAm profiles of a single tissue or cell type collected across childhood would be needed to disentangle this issue further. Unfortunately, such a dataset is not currently available as most cohorts have generated DNAm profiles from peripheral blood and cord blood^63^. Analysis of these complex tissues has nevertheless yielded many valuable insights. Second, since DNAm at 9 years was measured only in Generation R and at 17 years only in ALSPAC, DNAm differences from 9 to 17 may be to some extent driven by batch effects or cohort differences. This may explain some of the inter-individual differences in slope changes at 9 towards 17 years. However, the high level of agreement in both stability and change among the corresponding time points of the two cohorts is reassuring. Moreover, it is not entirely surprising that inter-individual variation in directionality of change was higher for the largest age interval. This interval, furthermore, encompasses the period of adolescent development, a time in which many inter-individual phenotypic differences arise. Finally, it should be noted that the current study only included children of European ancestry. Considerable DNAm differences have been found between populations^76–78^, but research on age-associated DNAm differences is scarce. One study^79^ reported evidence for overlap in age-associated CpGs in two African populations with studies on European-ancestry populations, but more research is needed to map the generalizability of longitudinal DNAm changes among different populations.

In conclusion, in the first comprehensive CpG-by-CpG characterization of DNAm from birth to late adolescence, we found that DNAm at more than half of the studied CpG sites changes consistently between individuals, and that considerable inter-individual variation in change exists. Further, characteristics such as child sex, CpG location, and environmental and disease traits have distinct associations with patterns of DNAm change. Further analysis of these patterns is made readily available at http://epidelta.mrcieu.ac.uk/, which we hope can be used in future studies to test developmental hypotheses that promote our understanding of the developmental nature of DNAm, its role in gene functioning, and the associated biological pathways leading to health and disease.

## Methods

### Setting

Data were obtained from two population-based prospective birth cohorts, the Dutch Generation R Study (Generation R) and the British Avon Longitudinal Study of Parents and Children (ALSPAC). Pregnant women residing in the study area of Rotterdam, the Netherlands, with an expected delivery date between April 2002 and January 2006 were invited to enroll in Generation R. A more extensive description of the study can be found elsewhere^80^. The Generation R Study is conducted in accordance with the World Medical Association Declaration of Helsinki and has been approved by the Medical Ethics Committee of the Erasmus Medical Center, Rotterdam. Informed consent was obtained for all participants.

Pregnant women residing in the study area of former county Avon, United Kingdom, with an expected delivery date between April 1991 and December 1992 were invited to enroll in the ALSPAC study. Detailed information on the study design can be found elsewhere^3,81^. The ALSPAC website contains details of all available data through a fully searchable data dictionary and variable search tool (http://www.bristol.ac.uk/alspac/researchers/our-data/). Ethical approval for the study was obtained from the ALSPAC Ethics and Law Committee and the Local Research Ethics Committees. Consent for biological samples has been collected in accordance with the Human Tissue Act (2004). Informed consent for the use of data collected via questionnaires and clinics was obtained from participants following the recommendations of the ALSPAC Ethics and Law Committee at the time.

### Study Population

In the Generation R Study, 9,778 pregnant mothers had 9,749 live-born children. For a subsample of 1,414 children DNAm data was collected at birth and/or 6 years and/or 10 years of age. This subsample consisted of participants with parents born in the Netherlands (European ancestry^82^ confirmed for all children with genetic data available (95.4%)). Fifteen sibling pairs were present in the dataset. From each pair one sibling with the lowest number of DNAm measurements, or otherwise randomly, was excluded, resulting in a sample with 1,399 children (with 2,333 DNAm samples; see below).

In the ALSPAC study, 15,247 pregnant mothers gave birth to 14,973 live-born children. DNAm at birth and/or 7 years and/or 17 years was available for a subsample of 1,003 children as part of the Accessible Resource for Integrated Epigenomic Studies (ARIES) study^83^. From this sample, 48 children with non-European ancestry as based on genetic principle component analysis and 6 children with missing data on gestational age were excluded, resulting in a sample of 949 children with DNAm data (with 2,686 DNAm samples; see below).

### DNA methylation

Cord blood was drawn after birth for both cohorts, and peripheral blood was drawn at a mean age of 6.0 (*SD*=0.47) and 9.8 (*SD*=0.3) years for Generation R, and 7.5 (*SD*=0.2) and 17.1 (*SD*=1.0) years for ALSPAC. Both cohorts made use of the EZ-96 DNAm kit (shallow) (Zymo Research Corporation, Irvine, USA) to perform bisulfite conversion on the extracted leukocytic DNA. Samples were further processed with the Illumina Infinium HumanMethylation450 BeadChip (Illumina Inc., San Diego, USA) to analyze DNAm.

In Generation R, quality control was performed on all 2,467 available DNAm samples with the CPACOR workflow^84^. Arrays with observed technical problems such as failed bisulfite conversion, hybridization or extension, as well as arrays with a mismatch between sex of the proband and sex determined by the chromosome X and Y probe intensities were removed from subsequent analyses. Additionally, only arrays with a call rate >95% per sample were processed further, resulting in 2,355 samples, 22 of which belonged to half of an excluded sibling pair, hence 2,333 samples were carried forward into normalization.

In ALSPAC, quality control was performed on 6,057 samples (3,286 belonging to children, 2,771 to their mothers), using the *meffil* package^85^ in R version 3.4.3^86^. After removing samples with mismatched genotypes, mismatched sex, incorrect relatedness, low concordance with samples collected at other time points, extreme dye bias, and poor probe detection, 5,337 samples remained, 2,845 of which belonging to children, used in the current study.

To minimize cohort effects as much as possible, we normalized both cohorts together as a single dataset. Functional normalization (10 control probe principal components, slide included as a random effect) was performed with the *meffil* package in R^85^. Normalization took place on the combined Generation R and ALSPAC set comprising a total of 5,178 samples for a total of 485,512 CpGs. One-hundred and fifty-nine ALSPAC samples belonging to non-European children or children with missing data on gestational age were excluded, leading to a final ALSPAC set of 2,686 samples (for 949 children). Together with 2,333 samples for Generation R (of 1,399 children) they formed a combined set of 5,019 samples (of 2,348 children.)

Analyses were restricted to 473,864 autosomal CpGs. DNAm levels were operationalized as beta values (β values), representing the ratio of methylated signal relative to the sum of methylated and unmethylated signal measured per CpG.

### Covariates

Sample plate number (*N*=29 in Generation R and *N*=31 in ALSPAC), was used to correct for batch effects, which was added as a random variable in the model (see below). White blood cell (WBC) composition was estimated with the reference-based Bakulski method^87^ for cord blood and Houseman method^88^ for peripheral blood (Supplemental Table 11). Nucleated red blood cells were not further analyzed due to its specificity to cord blood, leaving CD4+ T-lymphocytes, CD8+ T-lymphocytes, natural killer cells, B-lymphocytes, monocytes, and granulocytes. Other covariates included gestational age in weeks, sex of the child, and cohort.

### Statistical analyses

#### Step 1: Assessing cross-cohort comparability in DNA methylation stability

To ascertain comparability amongst the two cohorts we compared within-cohort DNAm stability between the time points that were present in both cohorts – i.e. birth and 6/7 years (Generation R/ALSPAC, respectively).

Longitudinal stability per CpG within each cohort was assessed by studying estimates of concordance and change. For concordance, DNAm data was first residualized within each cohort for all variables present in the longitudinal models except the ‘cohort’ variable, in order to remove between-cohort differences due to other covariates. Concordance was then measured both with Spearman correlation (data at most CpGs is not normally distributed) as a measure of relative concordance, and with intra-class correlations as a measure of absolute concordance (children with data for both time points: *n* Generation R=476, *n* ALSPAC=826). Longitudinal change from birth to 6/7 years was assessed by studying the estimates of the change in DNAm per year by applying Model 1 (see below) within each cohort (children with data for at least one of the two time-points: *n* Generation R=1,394, *n* ALSPAC=944).

In a second step, cross-cohort comparability was assessed with Spearman (ρ) correlation of concordance estimates of the CpGs of each cohort (which were not normally distributed) and Pearson correlations (*r*) amongst the change estimates of the CpGs of each cohort (which were normally distributed).

#### Step 2: Longitudinal modelling of DNA methylation using combined Generation R and ALSPAC data

The combined Generation R and ALSPAC dataset had four time points of collection (birth, age 6/ 7 years, 10 years, and 17 years). We fit three linear mixed models to CpG site DNAm across the genome to identify (i) linear change over time (Model 1); (ii) nonlinear change over time (Model 2); and (iii) sex differences in change over time (Model 3). Both fixed and random effects were examined to allow for inter-individual variation in DNAm patterns over time. The models are described in detail below.

##### Model 1: Linear change

This model was applied to identify CpGs that show an overall change in DNAm from birth to 18 years (i.e. fixed age effect), as well as CpGs with inter-individual differences in change during that time (i.e. random age effect). The Model 1 is defined as:

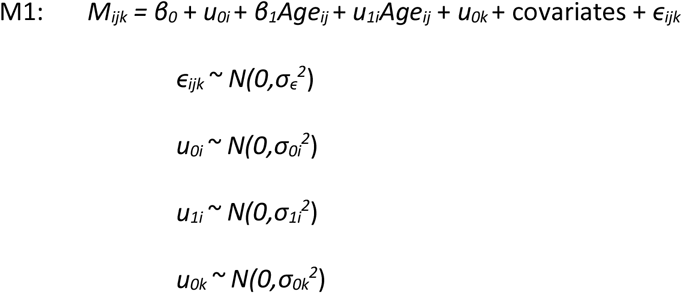

Here, participants are denoted by *i*, time points by *j*, and sample plates by *k*. *M* denotes DNAm level, *β*_*0*_ fixed intercept, *u*_*0i*_ random intercept, *β*_*1*_ fixed age coefficient, *u*_*1i*_ random age coefficient, *u*_*0k*_ random intercept for sample plate. Hence, *β*_*1*_ represents the average change in DNAm per one year. Variability in this change amongst individuals was captured with *u_1i_.* To avoid problems with model identification, the random slope of age was uncorrelated to the random intercept (i.e. a diagonal random effects matrix was used).

##### Model 2: Nonlinear change

To identify nonlinear changes in DNAm, we extended Model 1 to allow slope changes at ages 6 and 9^30,31^:

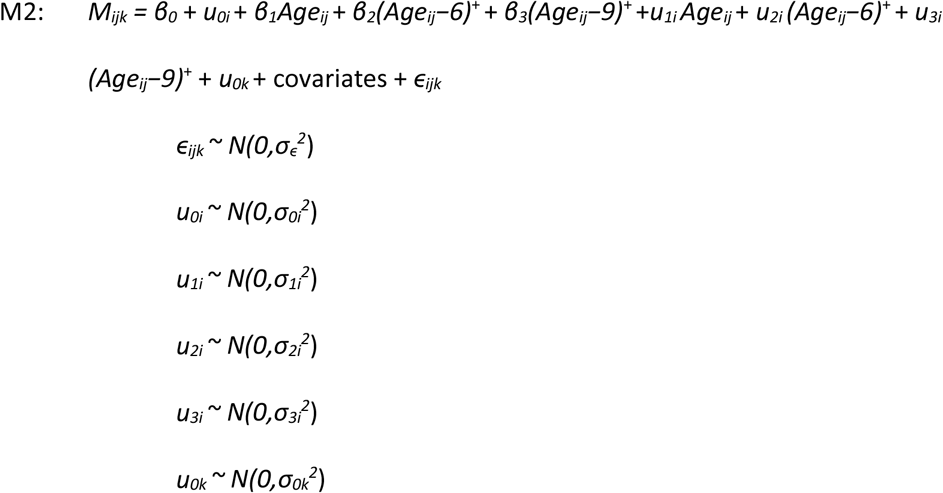

Where a^+^ = a if a>0 and 0 otherwise, so that *β*_*2*_ represents the average change in DNAm per year from 6 years of age onward, after accounting for the change per year from birth onward, as denoted by *β*_*1*_. Likewise, *β*_*3*_represents the average change in DNAm per year from 9 years of age onward, after accounting for the change per year from 6 years of age onward. Hence, with those variables we are able to detect slope changes at 6 and 9 years old. These slope changes were used to identify different types of nonlinear patterns. With *u*_*2i*_ and *u*_*3i*_ the inter-individual variation in slope changes at 6 and 9 years were captured, respectively. General linear hypothesis testing^89^ was applied to our fitted models to determine if there were changes in DNAm per year from 6-9 years and from 9-18 years.

##### Model 3: Sex differences in change

To identify CpGs for which DNAm changes differently over time for boys and girls, we applied the following model:

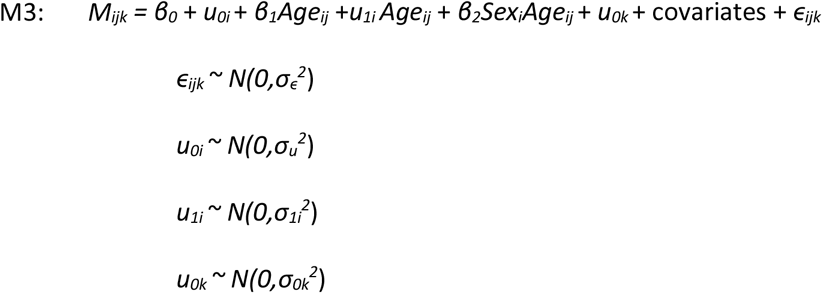

Here, *Sex*_*i*_ denotes the sex of child *i*. Both main and interaction effects for sex were studied.

The three mixed models were fitted using maximum likelihood estimation in *R* with the *lme4* package^90^. Continuous covariates (WBCs, gestational age) were z-score standardized. Random slopes were kept uncorrelated with random intercepts and the NLopt optimizer was used, enabling us to improve computational speed compared to the default settings. *P*-values for the fixed effects were computed with a z-test. *P*-values for random slopes of the Age effects were obtained by refitting the model without the random slope and comparing the fit estimates of the two models with a likelihood ratio test. Within each model, *P*-value thresholds were Bonferroni-corrected for the number of tested CpGs (i.e. to *P*<1×10^−07^).

#### Step 3: Functional characterization of probes with comparable patterns of change

To interpret the functionality of the age-related DNAm patterns from the three models, CpG sites adhering to 8 different age-related patterns (M1 linear change and inter-individual variation in linear change, M2 nonlinear trajectories, and inter-individual variation in change from birth, in slope change at 6 years, and in slope change at 9 years, and M3 stable sex differences and sex differences in DNAm change) were tested for enrichment in:

i. gene-relative genomic regions (TSS1500, TSS200, 5’UTR, 1st exon, gene body, 3’UTR, and intergenic regions^64^),
ii. CpG island-relative genomic regions (N shelf, N shore, CpG island, S shore, S shelf, and open sea regions^64^) as indicated by the Illumina HumanMethylation450 v1.2 Manifest (Illumina Inc., San Diego, USA), and
iii. enhancer elements as those expressed in whole blood, peripheral blood mononuclear cells, natural killer cells, CD4+ T cells, CD8+ T cells, monocytes, neutrophils, eosinophils or B cells^91^,

Altogether, these encompassed 14 enrichment analyses for 8 variables. Enrichment was tested using χ^2^-tests of unequal proportions. The enrichment *P*-value threshold was Bonferroni-corrected for multiple tests (i.e. *P*<4.46×10^−04^ for 8×14=122 tests). Second, we tested enrichment of Gene Ontology (GO) categories for genes linked to CpG sites surviving adjustment for multiple tests (*P*<1×10^−07^) for each of the main variables of interest. This analysis was adjusted for gene size and pruned for near-identical terms (see elsewhere for a full description^92^). For completeness, terms with nominal *P*<0.05 were reported. Last, we tested enrichment of age-related DNAm trajectories (11 different age-related patterns: M1 decreasing, increasing, and inter-individual variation in linear change, M2 Positive-Neutral, Negative-Neutral, other nonlinear, inter-individual variation in change from birth, in slope change at 6 years, and in slope change at 9 years, and M3 stable sex differences and sex differences in DNAm change) in EWASs on age, prenatal smoking, smoking, cardiovascular-associated traits, C-reactive protein, allergies, educational attainment, and cellular heterogeneity. EWAS summary statistics were retrieved from the EWAS Catalog (http://www.ewascatalog.org/) and studies were included when performed with the 450K array in peripheral or cord blood, resulting in 21 EWASs. Enrichment was tested with Fisher’s exact tests, the enrichment *P*-value threshold was Bonferroni-corrected for multiple tests (i.e. *P*<1.37×10^−04^ for 11×33=363 tests).

## Data access

Data from the Generation R Study are available upon request, subject to local rules and regulations. ALSPAC data access is through a system of managed open access. The ALSPAC access policy (http://www.bristol.ac.uk/media-library/sites/alspac/documents/researchers/data-access/ALSPAC_Access_Policy.pdf) describes the process of accessing the data and samples in detail, and outlines the costs associated with doing so. The results per CpG will be made available at http://epidelta.mrcieu.ac.uk.

## Acknowledgements

The Generation R Study is conducted by the Erasmus Medical Center in close collaboration with the Faculty of Social Sciences of the Erasmus University Rotterdam, the Municipal Health Service Rotterdam area, Rotterdam, the Rotterdam Homecare Foundation, Rotterdam and the Stichting Trombosedienst & Artsenlaboratorium Rijnmond (STAR-MDC), Rotterdam. We gratefully acknowledge the contribution of children and parents, general practitioners, hospitals, midwives and pharmacies in Rotterdam. The study protocol was approved by the Medical Ethical Committee of the Erasmus Medical Centre, Rotterdam. Written informed consent was obtained for all participants. The generation and management of the Illumina 450K methylation array data (EWAS data) for the Generation R Study was executed by the Human Genotyping Facility of the Genetic Laboratory of the Department of Internal Medicine, Erasmus MC, the Netherlands. We thank Mr Michael Verbiest, Ms Mila Jhamai, Ms Sarah Higgins, Mr Marijn Verkerk and Dr. Lisette Stolk for their help in creating the EWAS database. We thank Dr A. Teumer for his work on the quality control and normalization scripts. The general design of the Generation R Study is made possible by financial support from the Erasmus Medical Center, Rotterdam, the Erasmus University Rotterdam, the Netherlands Organization for Health Research and Development and the Ministry of Health, Welfare and Sport. The EWAS data was funded by a grant from the Netherlands Genomics Initiative (NGI)/Netherlands Organisation for Scientific Research (NWO) Netherlands Consortium for Healthy Aging (NCHA; project nr. 050-060-810), by funds from the Genetic Laboratory of the Department of Internal Medicine, Erasmus MC, and by a grant from the National Institute of Child and Human Development (R01HD068437). This project further received funding from the European Union’s Horizon 2020 research and innovation programme (733206, LifeCycle; 848158, EarlyCause; 874739, LongITools) and from the European Joint Programming Initiative “A Healthy Diet for a Healthy Life” (JPI HDHL, NutriPROGRAM project, ZonMw the Netherlands no.529051022 and PREcisE project ZonMw the Netherlands no.529051023).

We are extremely grateful to all the families who took part in the ALSPAC study, the midwives for their help in recruiting them, and the whole ALSPAC team, which includes interviewers, computer and laboratory technicians, clerical workers, research scientists, volunteers, managers, receptionists and nurses. The UK Medical Research Council and Wellcome (grant number 102215/2/13/2) and the University of Bristol provide core support for ALSPAC. A comprehensive list of grants funding is available on the ALSPAC website (http://www.bristol.ac.uk/alspac/external/documents/grant-acknowledgements.pdf). This publication is the work of the authors and Dr Suderman will serve as guarantor for the ALSPAC-related contents of this paper. Analysis of the ALSPAC data was funded by UK Economic & Social Research Council grant (grant number ES/N000498/1). ARIES was funded by the BBSRC (BBI025751/1 and BB/I025263/1). Supplementary funding to generate DNA methylation data which are (or will be) included in ARIES has been obtained from the MRC, ESRC, NIH and other sources. ARIES is maintained under the auspices of the MRC Integrative Epidemiology Unit at the University of Bristol (grant numbers MC_UU_00011/4 and MC_UU_00011/5).

AN and HT are supported by a grant of the Dutch Ministry of Education, Culture, and Science and the Netherlands Organization for Scientific Research (NWO grant No. 024.001.003, Consortium on Individual Development). HT was also supported by a grant from the Netherlands Organization for Scientific Research (NWO-grant 016.VICI.170.200). EW is funded by CLOSER, who is funded by ESRC and MRC (grant reference: ES/K000357/1). The funders took no role in the design, execution, analysis or interpretation of the data or in the writing up of the findings. www.closer.ac.uk. The work of CAMC has received funding from the European Union’s Horizon 2020 research and innovation programme under the Marie Skłodowska-Curie grant agreement No 707404. LCH is supported by ESRC/BBSRC Grant ES/N000382/1 for the Interpreting epigenetic signatures in studies of early life adversity project (Interstela Project). JR is supported by the Netherlands Organization for Scientific Research (NWO ZonMw VENI, grant no 91618147). VWVJ received funding from the European Research Council (ERC-2014-CoG-648916). MJB-K is supported by the European Research Council (ERC AdG) and by the Dutch Ministry of Education, Culture, and Science and the Netherlands Organization for Scientific Research (NWO grant No. 024.001.003, Consortium on Individual Development). MHvIJ is supported by the Dutch Ministry of Education, Culture, and Science and the Netherlands Organization for Scientific Research (NWO grant No. 024.001.003, Consortium on Individual Development) and by a Spinoza Award of the Netherlands Organization for Scientific Research.

## Author contributions

RHM, AN, EW, CAMC, LCH, JR, BTH, MJB-K, HT, MvIJ, CLR, and MS designed the study; RHM, AN, JR, TRG, JFF, HT, VWJ and CLR acquired the data; RHM and MS analyzed the data; RHM, AN, EW, CAMC, LCH, AJS, BTH, JFF, MJB-K, MvIJ, and MS interpreted the data; RHM drafted the work and all authors substantively revised it. All authors have approved the submitted version. All authors have agreed both to be personally accountable for the author’s own contributions and to ensure that questions related to the accuracy or integrity of any part of the work are appropriately investigated, resolved, and the resolution documented in the literature. The funders took no role in the design, execution, analysis or interpretation of the data or in the writing up of the findings.

