## Supplementary Material for "Epigenome-wide change and variation in DNA methylation from birth to late adolescence"

Supplementary Table 1. Cohort characteristics

|  | | **Generation R**  **(N = 1399)** | **ALSPAC**  **(N = 949)** |
| --- | --- | --- | --- |
| **Child characteristics** | |  |  |
| Sex (% boys) | | 50.8 | 48.7 |
| Gestational age (weeks) | | 40.1 (1.5) | 39.6 (1.5) |
| Age (years) | |  |  |
| @6 or 7 years, n = 488 / 970 | | 6.0 (0.4) | 7.5 (0.1) |
| @10 years, n = 463 | | 9.8 (0.3) |  |
| @17 years, n = 970 | |  | 17.1 (1.0) |
| Birth weight (grams) | | 3546 (510) | 3489 (490) |
| BMI | |  |  |
| @6 or 7 years, n = 488 / 970 | | 15.9 (1.3) | 16.2 (2.0) |
| @10 years, n = 463 | | 17.1 (2.0) |  |
| @17 years, n = 970 | |  | 22.5 (3.8) |
| **Mother characteristics** | |  |  |
| Maternal age at birth (years) | | 32.2 (4.2) | 30.0 (4.4) |
| BMI early pregnancy | | 24.2 (4.0) | 22.8 (3.7) |
| Education level (%) |  |  |  |
| low |  | 11.2 | 8.6 |
| medium |  | 23.5 | 41.1 |
| high |  | 65.3 | 50.3 |
| Prenatal smoking (% sustained) | | 13.3 | 10.2 |

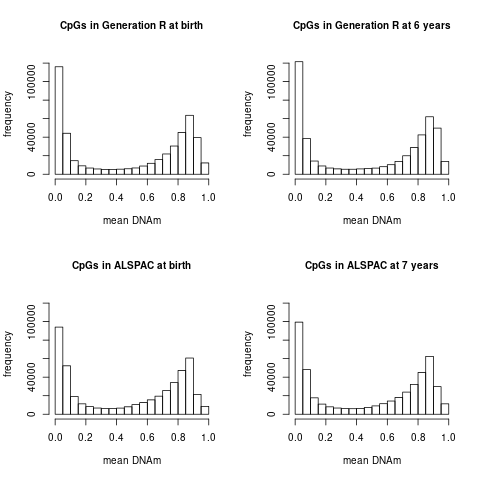

Supplementary Figure 1. Distribution of CpG mean methylation levels in (a) Generation R at birth; (b) ALSPAC at birth; (c) Generation R 6 years; (d) ALSPAC at 7 years

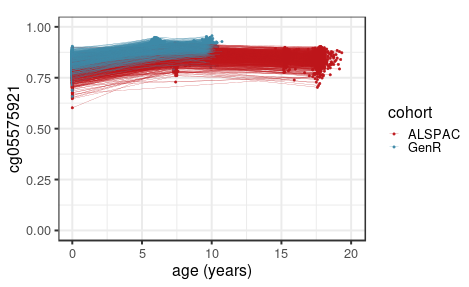

Supplementary Figure 2. Longitudinal DNA methylation of *AHRR* cg05575921 as predicted by Model 2.

Supplementary Tables 3A-H. CpG characteristics per gene region compared to CpGs at all other regions

Supplementary Table 3A. M1 linear DNAm change

|  | CpGs in area _ | other CpGs _ |  |  |  |
| --- | --- | --- | --- | --- | --- |
|  | % null/negative/  positive | % null/negative/  positive | χ^2^ | df | *P* |
| TSS1500 | 48.63/31.78/19.59 | 48.41/36.30/15.29 | 1164.52 | 2 | 1.34x10^-253^ |
| TSS200 | 52.79/28.19/19.02 | 47.82/36.59/15.60 | 1694.99 | 2 | <9.88x10^-324^ |
| 5'UTR | 50.50/30.06/19.44 | 48.13/36.36/15.51 | 1200.84 | 2 | 1.7 x10^-261^ |
| 1st exon | 51.63/25.35/23.02 | 48.17/36.4/15.43 | 2527.50 | 2 | <9.88x10^-324^ |
| Gene body | 48.31/38.75/12.94 | 48.53/33.67/17.80 | 2420.25 | 2 | <9.88x10^-324^ |
| 3'UTR | 48.03/42.77/9.20 | 48.47/35.21/16.32 | 880.58 | 2 | 6.08x10^-192^ |
| Intergenic | 45.71/38.73/15.56 | 49.35/34.46/16.19 | 718.15 | 2 | 1.14x10^-156^ |

Supplementary Table 3B. M1 inter-individual variation in DNAm change

|  | CpGs in area_ | other CpGs_ |  |  |  |
| --- | --- | --- | --- | --- | --- |
|  | % no/yes | % no/yes | χ^2^ | df | *P* |
| TSS1500 | 77.10/22.90 | 71.71/28.29 | 984.90 | 1 | 3.43x10^-216^ |
| TSS200 | 77.76/22.24 | 71.89/28.11 | 908.37 | 1 | 1.49x10^-119^ |
| 5'UTR | 76.11/23.89 | 72.10/27.9 | 442.63 | 1 | 2.90x10^-98^ |
| 1st exon | 76.33/23.67 | 72.31/27.69 | 281.83 | 1 | 3.00x10^-63^ |
| Gene body | 71.14/28.86 | 73.49/26.51 | 305.25 | 1 | 2.36x10^-68^ |
| 3'UTR | 72.79/27.21 | 72.63/27.37 | 0.24 | 1 | 6.24x10^-01^ |
| Intergenic | 69.42/30.58 | 73.70/26.3 | 814.75 | 1 | 3.35x10^-179^ |

Supplementary Table 3C. M2 nonlinear DNAm change

|  | CpGs in area _ | other CpGs _ |  |  |  |
| --- | --- | --- | --- | --- | --- |
|  | % linear or null/  Positive-Neutral/  Negative-Neutral/  other nonlinear | % linear or null/  Positive-Neutral/  Negative-Neutral/  other nonlinear | χ^2^ | df | *P* |
| TSS1500 | 88.55/4.70/3.77/2.98 | 89.11/4.84/3.00/3.05 | 132.58 | 3 | 1.50x10^-28^ |
| TSS200 | 92.85/3.14/1.38/2.63 | 88.46/5.06/3.39/3.09 | 1227.78 | 3 | 6.90x10^-266^ |
| 5'UTR | 89.73/4.13/3.06/3.08 | 88.91/4.92/3.15/3.03 | 75.95 | 3 | 2.27x10^-16^ |
| 1st exon | 92.11/3.21/1.56/3.12 | 88.75/4.95/3.27/3.03 | 589.94 | 3 | 1.53x10^-127^ |
| Gene body | 88.40/4.84/3.66/3.10 | 89.37/4.80/2.84/2.99 | 252.42 | 3 | 1.96x10^-54^ |
| 3'UTR | 88.91/4.64/3.40/3.05 | 89.02/4.82/3.12/3.03 | 5.89 | 3 | 1.17x10^-01^ |
| Intergenic | 88.12/5.82/2.92/3.14 | 89.31/4.48/3.21/3.00 | 371.72 | 3 | 2.96x10^-80^ |

Supplementary Table 3D. M2 inter-individual variation in DNAm change from birth

|  | CpGs in area_ | other CpGs_ |  |  |  |
| --- | --- | --- | --- | --- | --- |
|  | % no/yes | % no/yes | χ^2^ | df | *P* |
| TSS1500 | 96.99/3.01 | 96.58/3.42 | 35.70 | 3 | 2.30x10^-09^ |
| TSS200 | 95.98/4.02 | 96.75/3.25 | 95.82 | 3 | 1.26x10^-22^ |
| 5'UTR | 96.20/3.80 | 96.72/3.28 | 45.97 | 3 | 1.20x10^-11^ |
| 1st exon | 95.09/4.91 | 96.79/3.21 | 308.77 | 3 | 4.05x10^-69^ |
| Gene body | 97.04/2.96 | 96.43/3.57 | 128.26 | 3 | 9.82x10^-30^ |
| 3'UTR | 97.89/2.11 | 96.60/3.40 | 94.92 | 3 | 1.98x10^-22^ |
| Intergenic | 96.54/3.46 | 96.69/3.31 | 6.45 | 3 | 1.11x10^-02^ |

Supplementary Table 3E. M2 inter-individual variation in DNAm change from 6 years

|  | CpGs in area_ | other CpGs_ |  |  |  |
| --- | --- | --- | --- | --- | --- |
|  | % no/yes | % no/yes | χ^2^ | df | *P* |
| TSS1500 | 99.83/0.17 | 99.83/0.17 | 0.21 | 2 | 6.49x10^-01^ |
| TSS200 | 99.64/0.36 | 99.86/0.14 | 149.48 | 2 | 2.25x10^-34^ |
| 5'UTR | 99.72/0.28 | 99.85/0.15 | 56.08 | 2 | 6.96x10^-14^ |
| 1st exon | 99.47/0.53 | 99.86/0.14 | 310.00 | 2 | 2.18x10^-69^ |
| Gene body | 99.90/0.10 | 99.80/0.20 | 63.49 | 2 | 1.61x10^-15^ |
| 3'UTR | 99.94/0.06 | 99.83/0.17 | 13.93 | 2 | 1.90x10^-04^ |
| Intergenic | 99.9/0.10 | 99.81/0.19 | 43.23 | 2 | 4.88x10^-11^ |

Supplementary Table 3F. M2 inter-individual variation in DNAm change from 9 years

|  | CpGs in area_ | other CpGs_ |  |  |  |
| --- | --- | --- | --- | --- | --- |
|  | % no/yes | % no/yes | χ^2^ | df | *P* |
| TSS1500 | 93.34/6.66 | 91.52/8.48 | 296.39 | 2 | 2.01x10^-66^ |
| TSS200 | 91.47/8.53 | 91.89/8.11 | 12.44 | 2 | 4.21x10^-04^ |
| 5'UTR | 92.08/7.92 | 91.80/8.20 | 5.76 | 2 | 1.64x10^-02^ |
| 1st exon | 90.79/9.21 | 91.92/8.08 | 59.47 | 2 | 1.24x10^-14^ |
| Gene body | 91.18/8.82 | 92.21/7.79 | 156.20 | 2 | 7.64x10^-36^ |
| 3'UTR | 92.86/7.14 | 91.79/8.21 | 28.01 | 2 | 1.20x10^-07^ |
| Intergenic | 92.33/7.67 | 91.67/8.33 | 51.46 | 2 | 7.30x10^-13^ |

Supplementary Table 3G. M3 stable sex differences in DNAm

|  | CpGs in area_ | other CpGs_ |  |  |  |
| --- | --- | --- | --- | --- | --- |
|  | % no/yes | % no/yes | χ^2^ | df | *P* |
| TSS1500 | 94.49/5.51 | 95.33/4.67 | 544.43 | 2 | 5.99x10^-119^ |
| TSS200 | 96.66/3.34 | 94.97/5.03 | 400.83 | 2 | 9.14x10^-88^ |
| 5'UTR | 96.41/3.59 | 95.00/5.00 | 237.15 | 2 | 3.19x10^-52^ |
| 1st exon | 96.23/3.77 | 95.09/4.91 | 100.04 | 2 | 1.89x10^-22^ |
| Gene body | 96.12/3.88 | 94.65/5.35 | 755.26 | 2 | 9.93x10^-165^ |
| 3'UTR | 96.45/3.55 | 95.13/4.87 | 103.99 | 2 | 2.63x10^-23^ |
| Intergenic | 93.18/6.82 | 95.85/4.15 | 1377.23 | 2 | 8.68x10^-300^ |

Supplementary Table 3H. M3 sex differences in DNAm change

|  | CpGs in area_ | other CpGs_ |  |  |  |
| --- | --- | --- | --- | --- | --- |
|  | % no/yes | % no/yes | χ^2^ | df | *P* |
| TSS1500 | 99.64/0.36 | 99.62/0.38 | 7.67 | 2 | 2.16x10^-02^ |
| TSS200 | 99.73/0.27 | 99.61/0.39 | 25.52 | 2 | 2.88x10^-06^ |
| 5'UTR | 99.68/0.32 | 99.62/0.38 | 6.16 | 2 | 4.60x10^-02^ |
| 1st exon | 99.76/0.24 | 99.62/0.38 | 28.74 | 2 | 5.74x10^-07^ |
| Gene body | 99.65/0.35 | 99.62/0.38 | 7.59 | 2 | 2.25x10^-02^ |
| 3'UTR | 99.70/0.30 | 99.62/0.38 | 2.96 | 2 | 2.28x10^-01^ |
| Intergenic | 99.51/0.49 | 99.66/0.34 | 54.36 | 2 | 1.57x10^-12^ |

Supplementary Tables 4A-H. Chi-square of CpG characteristics for each CpG island region compared to CpG characteristics in all other regions

Supplementary Table 4A. M1 linear DNAm change

|  | CpGs in area _ | other CpGs _ |  |  |  |
| --- | --- | --- | --- | --- | --- |
|  | % null/negative/  positive | % null/negative/  positive | χ^2^ | df | *P* |
| N shelf | 50.09/42.08/7.83 | 48.41/36.3/15.29 | 1395.43 | 2 | 9.68x10^-304^ |
| N shore | 49.11/31.25/19.64 | 47.82/36.59/15.60 | 940.13 | 2 | 7.13x10^-205^ |
| Island | 51.82/23.00/25.18 | 48.13/36.36/15.51 | 20798.58 | 2 | <9.88x10^-324^ |
| S shore | 49.26/31.66/19.08 | 48.17/36.40/15.43 | 540.32 | 2 | 4.68x10^-118^ |
| S shelf | 50.09/42.08/7.83 | 48.53/33.67/17.80 | 1344.60 | 2 | 1.05x10^-292^ |
| Open sea | 44.67/46.91/8.42 | 48.47/35.21/16.32 | 20603.72 | 2 | <9.88x10^-324^ |

Supplementary Table 4B. M1 inter-individual variation in DNAm change

|  | CpGs in area_ | other CpGs_ |  |  |  |
| --- | --- | --- | --- | --- | --- |
|  | % no/yes | % no/yes | χ^2^ | df | *P* |
| N shelf | 75.05/24.95 | 71.71/28.29 | 74.74 | 1 | 5.38x10^-18^ |
| N shore | 73.98/26.02 | 71.89/28.11 | 63.72 | 1 | 1.44x10^-15^ |
| Island | 71.40/28.60 | 72.10/27.90 | 160.31 | 1 | 9.69x10^-37^ |
| S shore | 74.26/25.74 | 72.31/27.69 | 70.67 | 1 | 4.23x10^-17^ |
| S shelf | 75.05/24.95 | 73.49/26.51 | 87.17 | 1 | 9.97x10^-21^ |
| Open sea | 72.06/27.94 | 72.63/27.37 | 44.92 | 1 | 2.05x10^-11^ |

Supplementary Table 4C. M2 nonlinear DNAm change

|  | CpGs in area _ | other CpGs _ |  |  |  |
| --- | --- | --- | --- | --- | --- |
|  | % linear or null/  Positive-Neutral/  Negative-Neutral/  other nonlinear | % linear or null/  Positive-Neutral/  Negative-Neutral/  other nonlinear | χ^2^ | df | *P* |
| N shelf | 88.51/5.04/3.89/2.56 | 89.04/4.80/3.10/3.06 | 68.65 | 3 | 8.31x10^-15^ |
| N shore | 86.10/6.79/4.50/2.62 | 89.45/4.52/2.93/3.10 | 1090.63 | 3 | 3.94x10^-236^ |
| Island | 93.10/3.25/0.79/2.87 | 87.20/5.51/4.18/3.11 | 5182.32 | 3 | <9.88x10^-324^ |
| S shore | 85.75/6.86/4.73/2.66 | 89.38/4.58/2.96/3.08 | 984.09 | 3 | 5.08x10^-213^ |
| S shelf | 88.51/5.04/3.89/2.56 | 89.04/4.80/3.10/3.06 | 49.85 | 3 | 8.58x10^-11^ |
| Open sea | 87.56/4.85/4.02/3.57 | 89.85/4.79/2.63/2.73 | 992.66 | 3 | 7.03x10^-215^ |

Supplementary Table 4D. M2 inter-individual variation in DNAm change from birth

|  | CpGs in area_ | other CpGs_ |  |  |  |
| --- | --- | --- | --- | --- | --- |
|  | % no/yes | % no/yes | χ^2^ | df | *P* |
| N shelf | 98.09/1.91 | 96.58/3.42 | 163.36 | 3 | 2.09x10^-37^ |
| N shore | 96.90/3.10 | 96.75/3.25 | 13.13 | 3 | 2.91x10^-04^ |
| Island | 94.67/5.33 | 96.72/3.28 | 2544.09 | 3 | <9.88x10^-324^ |
| S shore | 96.79/3.21 | 96.79/3.21 | 3.25 | 3 | 7.15x10^-02^ |
| S shelf | 98.09/1.91 | 96.43/3.57 | 197.61 | 3 | 6.94x10^-45^ |
| Open sea | 97.78/2.22 | 96.60/3.40 | 1070.38 | 3 | 9.06x10^-235^ |

Supplementary Table 4E. M2 inter-individual variation in DNAm change from 6 years

|  | CpGs in area_ | other CpGs_ |  |  |  |
| --- | --- | --- | --- | --- | --- |
|  | % no/yes | % no/yes | χ^2^ | df | *P* |
| N shelf | 99.97/0.03 | 99.83/0.17 | 26.83 | 2 | 2.22x10^-07^ |
| N shore | 99.94/0.06 | 99.86/0.14 | 44.75 | 2 | 2.24x10^-11^ |
| Island | 99.55/0.45 | 99.85/0.15 | 1027.50 | 2 | 1.89x10^-225^ |
| S shore | 99.93/0.07 | 99.86/0.14 | 27.85 | 2 | 1.31x10^-07^ |
| S shelf | 99.97/0.03 | 99.80/0.20 | 24.15 | 2 | 8.93x10^-07^ |
| Open sea | 99.97/0.03 | 99.83/0.17 | 328.18 | 2 | 2.40x10^-73^ |

Supplementary Table 4F. M2 inter-individual variation in DNAm change from 9 years

|  | CpGs in area_ | other CpGs_ |  |  |  |
| --- | --- | --- | --- | --- | --- |
|  | % no/yes | % no/yes | χ^2^ | df | *P* |
| N shelf | 94.58/5.42 | 91.52/8.48 | 255.42 | 2 | 1.71x10^-57^ |
| N shore | 92.96/7.04 | 91.89/8.11 | 118.35 | 2 | 1.45x10^-27^ |
| Island | 88.40/11.60 | 91.80/8.20 | 3305.19 | 2 | <9.88x10^-324^ |
| S shore | 92.97/7.03 | 91.92/8.08 | 91.09 | 2 | 1.37x10^-21^ |
| S shelf | 94.58/5.42 | 92.21/7.79 | 244.06 | 2 | 5.13x10^-55^ |
| Open sea | 93.27/6.73 | 91.79/8.21 | 753.18 | 2 | 8.18x10^-166^ |

Supplementary Table 4G. M3 stable sex differences in DNAm

|  | CpGs in area_ | other CpGs_ |  |  |  |
| --- | --- | --- | --- | --- | --- |
|  | % no/yes | % no/yes | χ^2^ | df | *P* |
| N shelf | 96.54/3.46 | 95.33/4.67 | 104.49 | 2 | 2.04x10^-23^ |
| N shore | 92.65/7.35 | 94.97/5.03 | 1214.08 | 2 | 2.32x10^-264^ |
| Island | 95.46/4.54 | 95.00/5.00 | 99.69 | 2 | 2.25x10^-22^ |
| S shore | 92.24/7.76 | 95.09/4.91 | 1379.57 | 2 | 2.69x10^-300^ |
| S shelf | 96.54/3.46 | 94.65/5.35 | 94.48 | 2 | 3.05x10^-21^ |
| Open sea | 96.31/3.69 | 95.13/4.87 | 1553.00 | 2 | <9.88x10^-324^ |

Supplementary Table 4H. M3 sex differences in DNAm change

|  | CpGs in area_ | other CpGs_ |  |  |  |
| --- | --- | --- | --- | --- | --- |
|  | % no/yes | % no/yes | χ^2^ | df | *P* |
| N shelf | 99.62/0.38 | 99.62/0.38 | 0.37 | 2 | 8.29x10^-01^ |
| N shore | 99.45/0.55 | 99.61/0.39 | 66.45 | 2 | 3.72x10^-15^ |
| Island | 99.74/0.26 | 99.62/0.38 | 89.21 | 2 | 4.26x10^-20^ |
| S shore | 99.36/0.64 | 99.62/0.38 | 115.80 | 2 | 7.17x10^-26^ |
| S shelf | 99.62/0.38 | 99.62/0.38 | 4.16 | 2 | 1.25x10^-01^ |
| Open sea | 99.68/0.32 | 99.62/0.38 | 21.98 | 2 | 1.68x10^-05^ |

Supplementary Tables 5. CpGs inside and outside enhancer regions

|  | % enhancer CpGs | % other CpGs | χ^2^ | df | *P* |
| --- | --- | --- | --- | --- | --- |
| M1 linear DNAm change  null/negative/positive | 47.54/28.58/23.88 | 48.47/35.66/15.87 | 508.87 | 2 | 3.16x10^-111^ |
| M1 inter-individual variation in DNAm change  no/yes | 72.55/27.45 | 72.64/27.36 | 0.03 | 1 | 8.70x10^-01^ |
| M2 nonlinear DNAm change  linear or null/Positive-Neutral/Negative-Neutral/other nonlinear | 78.65/9.85/8.21/3.29 | 89.23/4.71/3.03/3.03 | 1434.20 | 3 | 1.11x10^-310^ |
| M2 inter-individual variation in DNAm change from birth  no/yes | 94.41/5.59 | 96.7/3.30 | 150.83 | 1 | 1.13x10^-34^ |
| M2 inter-individual variation in DNAm change from 6 years  no/yes | 99.83/0.17 | 99.83/0.17 | 0.00 | 1 | 1.00 |
| M2 inter-individual variation in DNAm change from 9 years  no/yes | 92.28/7.72 | 91.82/8.18 | 2.55 | 1 | 1.10x10^-01^ |
| M3 stable DNAm sex differences  no/yes | 94.93/5.07 | 95.19/4.81 | 1.30 | 1 | 2.55x10^-01^ |
| M3 sex differences in DNAm change  no/yes | 99.74/0.26 | 99.62/0.38 | 2.95 | 1 | 8.58x10^-02^ |

Supplementary Table 9. CpGs part of Horvath age estimator

|  | % Horvath CpGs | % other CpGs | χ^2^ | df | *P* |
| --- | --- | --- | --- | --- | --- |
| M1 linear DNAm change  null/negative/positive | 37.11/29.75/33.14 | 48.46/35.52/16.02 | 77.22 | 2 | 1.71x10^-17^ |
| M1 inter-individual variation in DNAm change  no/yes | 74.22/25.78 | 72.63/27.37 | 0.37 | 1 | 5.42x10^-01^ |
| M2 nonlinear DNAm change  linear or null/Positive-Neutral/Negative-Neutral/other nonlinear | 73.65/5.38/8.22/12.75 | 89.03/4.81/3.13/3.03 | 148.54 | 3 | 5.43x10^-32^ |
| M2 inter-individual variation in DNAm change from birth  no/yes | 95.75/4.25 | 96.65/3.35 | 0.63 | 1 | 4.28x10^-01^ |
| M2 inter-individual variation in DNAm change from 6 years  no/yes | 100/0 | 99.83/0.17 | 0.01 | 1 | 9.04x10^-01^ |
| M2 inter-individual variation in DNAm change from 9 years  no/yes | 93.2/6.8 | 91.83/8.17 | 0.71 | 1 | 4.00x10^-01^ |
| M3 stable DNAm sex differences  no/yes | 86.97/13.03 | 95.19/4.81 | 50.23 | 1 | 1.37x10^-12^ |
| M3 sex differences in DNAm change  no/yes | 98.58/1.42 | 99.63/0.37 | 7.73 | 1 | 5.44x10^-03^ |

Supplementary Table10. CpGs part of Hannum age estimator

|  | % Hannum CpGs | % other CpGs | χ^2^ | df | *P* |
| --- | --- | --- | --- | --- | --- |
| M1 linear DNAm change  null/negative/positive | 8.45/46.48/45.07 | 48.45/35.52/16.03 | 63.2 | 2 | 1.89x10^-14^ |
| M1 inter-individual variation in DNAm change  no/yes | 46.48/53.52 | 72.64/27.36 | 23.14 | 1 | 1.51x10^-06^ |
| M2 nonlinear DNAm change  linear or null/Positive-Neutral/Negative-Neutral/other nonlinear | 54.93/7.04/7.04/30.99 | 89.02/4.81/3.14/3.03 | 196.34 | 3 | 2.61x10^-42^ |
| M2 inter-individual variation in DNAm change from birth  no/yes | 83.10/16.90 | 96.65/3.35 | 36.22 | 1 | 1.76x10^-09^ |
| M2 inter-individual variation in DNAm change from 6 years  no/yes | 100/0 | 99.83/0.17 | 0 | 1 | 1.00 |
| M2 inter-individual variation in DNAm change from 9 years  no/yes | 71.83/28.17 | 91.84/8.16 | 35.27 | 1 | 2.87x10^-09^ |
| M3 stable DNAm sex differences  no/yes | 90.14/9.86 | 95.18/4.82 | 2.92 | 1 | 8.76x10^-01^ |
| M3 sex differences in DNAm change  no/yes | 100/0 | 99.63/0.37 | 0 | 1 | 1.00 |

Supplemental Table 11. Estimated white blood cell proportions

|  | *Generation R* | *ALSPAC* |
| --- | --- | --- |
|  | mean (SD) | mean (SD) |
| 0 years – Bakulski method |  |  |
| CD8T | 0.13 (0.05) | 0.09 (0.05) |
| CD4T | 0.16 (0.05) | 0.18 (0.06) |
| NK | 0.03 (0.03) | 0.01 (0.02) |
| Bcell | 0.10 (0.03) | 0.17 (0.04) |
| Mono | 0.09 (0.02) | 0.01 (0.02) |
| Gran | 0.41 (0.11) | 0.35 (0.10) |
| nRBC | 0.12 (0.07) | 0.20 (0.09) |
| 6 / 7 years – Houseman method |  |  |
| CD8T | 0.12 (0.04) | 0.04 (0.03) |
| CD4T | 0.17 (0.05) | 0.21 (0.05) |
| NK | 0.02 (0.02) | 0.19 (0.04) |
| Bcell | 0.13 (0.03) | 0.14 (0.03) |
| Mono | 0.06 (0.02) | 0.06 (0.03) |
| Gran | 0.53 (0.08) | 0.44 (0.08) |
| 10 years – Houseman method |  |  |
| CD8T | 0.12 (0.04) |  |
| CD4T | 0.18 (0.05) |  |
| NK | 0.03 (0.03) |  |
| Bcell | 0.11 (0.03) |  |
| Mono | 0.06 (0.02) |  |
| Gran | 0.52 (0.08) |  |
| 17 years – Houseman method |  |  |
| CD8T |  | 0.03 (0.03) |
| CD4T |  | 0.18 (0.05) |
| NK |  | 0.20 (0.05) |
| Bcell |  | 0.11 (0.03) |
| Mono |  | 0.07 (0.03) |
| Gran |  | 0.48 (0.09) |
